# Co-expression of MDM2 and CDK4 in transformed human mesenchymal stem cells induces dedifferentiated liposarcoma potency

**DOI:** 10.1101/293068

**Authors:** Yu Jin Kim, Minjung Sung, Dan Bi Yu, Mingi Kim, Ji-Young Song, Kyoung Song, Kyungsoo Jung, Yoon-La Choi

**Affiliations:** Laboratory of Cancer Genomics and Molecular Pathology, Samsung Medical Center, Sungkyunkwan University School of Medicine, Seoul, Korea; Department of Health Sciences and Technology, SAIHST, Sungkyunkwan University, Seoul, Korea; The Center for Companion Diagnostics, LOGONE Bio Convergence Research Foundation, Seoul, Korea; Department of Pathology and Translational Genomics, Samsung Medical Center, Sungkyunkwan University School of Medicine, Seoul, Korea

**Keywords:** Well-differentiated liposarcoma (WDLPS), Dedifferentiated liposarcoma (DDLPS), MDM2, CDK4, Transformed bone marrow stem cell (BMSC), Adipogenesis

## Abstract

Amplification and overexpression of MDM2 and CDK4 are well-known diagnostic criteria of well-differentiated liposarcoma (WDLPS)/dedifferentiated liposarcoma (DDLPS). Although it was reported that depletion of *MDM2* or *CDK4* decreased proliferation in DDLPS cell lines, it remains unclear whether MDM2 and CDK4 induce WDLPS/DDLPS tumorigenesis. We examined whether MDM2 and/or CDK4 produce WDLPS/DDLPS using transformed human bone marrow stem cells (BMSCs), 2H and 5H, with five oncogenic hits (overexpression of hTERT, TP53 degradation, RB inactivation, c-MYC stabilization, and overexpression of HRAS^v12^). *In vitro* functional experiments revealed that co-overexpression of MDM2 and CDK4 plays key roles in tumorigenesis by increasing cell growth and migration and inhibiting adipogenic differentiation potency compared to sole expression of MDM2 or CDK4. Using mouse xenograft models, we found that co-overexpression of MDM2 and CDK4 in 5H cells with five additional oncogenic mutations can develop proliferative DDLPS *in vivo*. Our results suggest that co-overexpression of MDM2 and CDK4 induces DDLPS tumour potency in transformed human BMSCs by accelerating cell growth and migration and blocking adipogenetic potential incooperation with multiple genetic factors.

## Introduction

Liposarcoma (LPS) is one of the most frequently occurring types of soft tissue sarcoma in adults (Dodd, 2012). According to histological characteristics, LPS consists of three categories: well-differentiated or dedifferentiated, myxoid/round cell, and pleomorphic LPSs (Dodd, 2012). Well-differentiated (WD) or dedifferentiated (DD)LPS is the most common subtype and contains supernumerary ring and/or giant rod chromosomes formed by amplification of chromosome 12q13-15,which contains several hundred genes, including *MDM2* and *CDK4* (Pedeutour *et al*., 1999). Amplification and overexpression of MDM2 and CDK4 are generally accepted as current WDLPS/DDLPS diagnostic criteria (Barretina *et al*., 2010; Italiano *et al*., 2009; Song *et al*., 2017).

MDM2 inhibits tumour suppressor p53 and is overexpressed in numerous cancers (Toledo and Wahl, 2006). MDM2 functions as aubiquitin ligase that targets p53 through the proteasomal degradation pathway, as well as participates in its own auto-degradation to prohibit MDM2 activity, inhibiting p53 during periods of cellular stress (Choi *et al*., 2018; Stommel and Wahl, 2004). CDK4 forms a complex with cyclin D, which then phosphorylates pRB. This prevents E2F from interacting with phosphorylated pRB, which causes the cell cycle to progress into the G1-S transition and increases cell proliferation (Day *et al*., 2009; Harbour *et al*., 1999; Shim *et al*., 2010). Knockdown of *MDM2* or *CDK4* decreased cell proliferation in DDLPS cells (Barretina *et al*., 2010). Despite their potency as driving factors, it remains unclear whether MDM2 and CDK4 induce WDLPS/DDLPS tumorigenesis.

It has been well-established that genetic manipulation of important tumour suppressor genes and oncogenes induces transformation of human BMSCs to various sarcomas *in vitro* or *in vivo* (Kirsch *et al*., 2007; Rodriguez *et al*., 2012; Rosland *et al*., 2009; Torsvik *et al*., 2010; Wang *et al*., 2005). However, studies have failed to model sarcomagenesis through the expression of fusion oncogenes in human mesenchymal stem cells (MSCs) (Cironi *et al*., 2009; Riggi *et al*., 2008). Recently, robust evidence showed that expression of the FUS-CHOP fusion protein may initiate myxoid liposarcoma in transformed human BMSCs (Rodriguez *et al*., 2013). Therefore, we examined whether MDM2 and/or CDK4 produce WDLPS/DDLPS using transformed human BMSCs.

Binh *et al*. found that immunoexpression of MDM2 or CDK4, and MDM2 and CDK4 were 100% (44/44), 90.9% (40/44),, and 90.9% (40/44) in WDLPS and 95.1% (58/61), 91.8% (56/61), and 90.2%(55/61) in DDLPS, respectively (Binh *et al*., 2005). Sirvent *et al*. reported that the immunopositivity of MDM2 and CDK4 were 76.5% (26/34) and 82.4% (28/34) in WDLPS and 100% (8/8) and 100% (8/8) in DDLPS, respectively (Sirvent *et al*., 2007). In our previous study, amplification of *MDM2* and *CDK4* were 100% (30/30) and 90% (27/30) in WDLPS and 100% (26/26) and 92.3% (24/26) in DDLPS, respectively (Lee *et al*., 2014). Based on the overexpression of both MDM2 and CDK4 in WDLPS/DDLPS, we examined whether co-overexpression of MDM2 and CDK4 drives WDLPS/DDLPS tumorigenesis using transformed human BMSCs.

## Results

### Transformed human BMSCs retain their stemness characteristics

To examine whether co-overexpression of MDM2 and CDK4 drives WDLPS/DDLPS tumorigenesis, we used two transformed BMSCs (2H and 5H cells) containing two to five different oncogenic mutations (Fig. 1A) (Rodriguez *et al.*, 2013). These oncogenic hits include the following: □) ectopic overexpression of human telomerase reverse transcriptase (hTERT), □) TP53 degradation by expression of E6 antigen of human papillomavirus-16 (HPV-16), □) RB family inactivation by expression of E7 antigen of HPV-16, □) c-MYC stabilization by expression of small T antigen of Simian virus 40 (SV40), and □) activation of mitogenic signal by expression of HRAS^v12^. The E6 antigen of HPV-16mediates TP53 degradation via the proteasomal degradation pathway as observed in MDM2. However, E6 and MDM2 are regulated through well-established distinct mechanisms (Camus *et al*., 2003; Camus *et al*., 2007). Therefore, none of the five oncogenic aspects were directly relevant to WDLPS/DDLPS.

**Figure 1.**
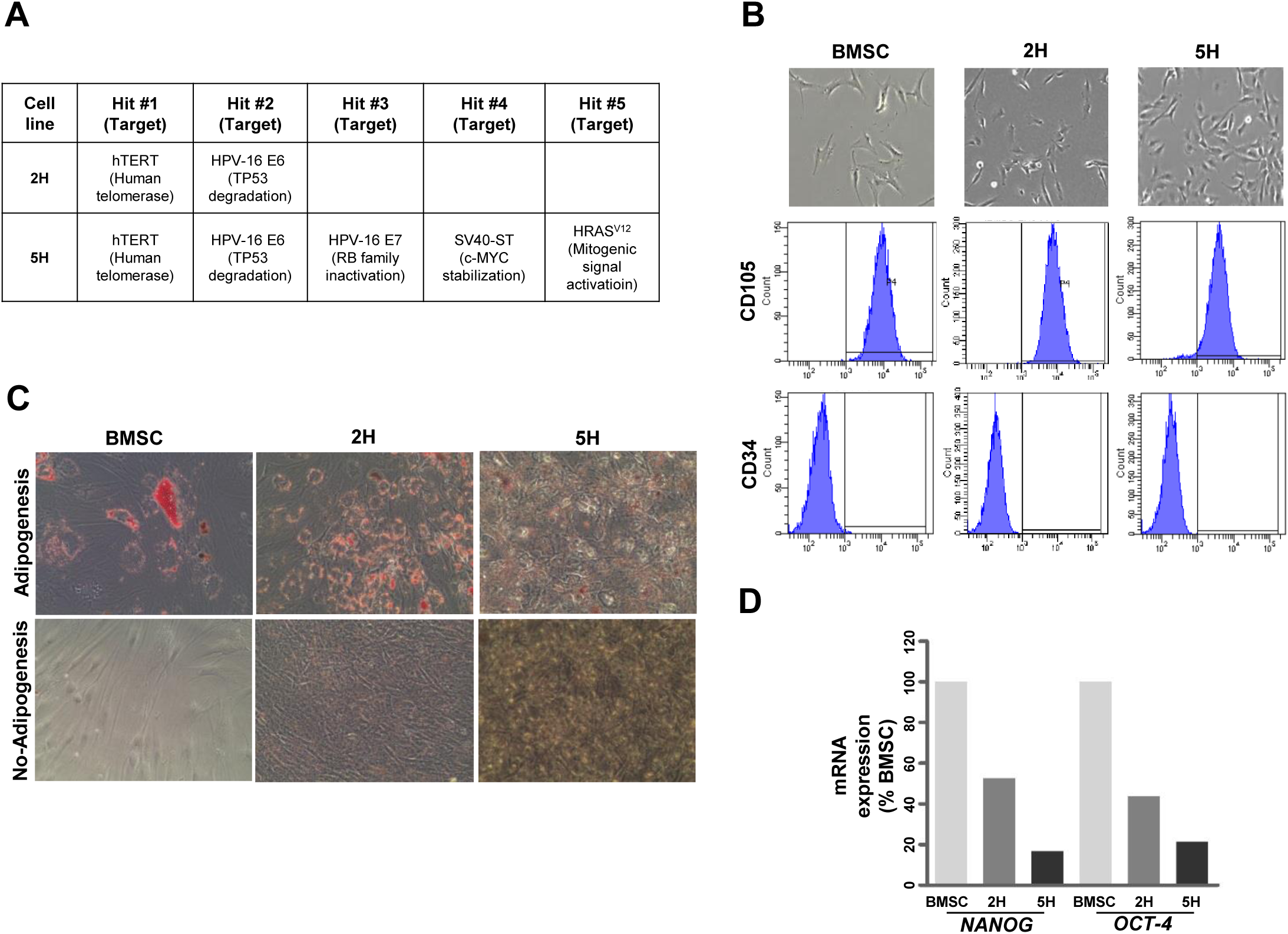
Transformed human BMSCs retain their stemness characteristics. (A) The oncogenic hits used in the 2H and 5H cells are indicated. (B) 2H and 5H cells underwent morphologic changes *in vitro*, adopting shorter and thicker appearances than BMSCs (upper panels). Expression of two cell surface makers (CD105 and CD34) was validated by fluorescence-activated cell sorting (FACS) analyses based on isotype-matched control antibodies (lower panels). (C) mRNA expression levels of *NANOG* and *OCT-4* were quantified using quantitative RT-PCR after induction adipogenesis. The percentage values were calculated based on their levels in BMSCs. (D) Ability to differentiate into adipocytes was monitored by Oil Red O staining.

We examined the characteristics of 2H and 5H cells and compared them to non-transformed BMSCs. The 2H and 5H cells expressed cell surface makers, such as CD105 (human mesenchymal stromal cell marker; positive) and CD34 (hematopoietic progenitor cell marker; negative), which is consistent with BMSCs (Fig. 1B). However, the morphology of 2H and 5H cells differed from that of BMSCs, which are characterized by spindle-shaped morphology that includes a large cell body with a long and thin tails. Both 2H and 5H cells were shorter and thicker than BMSCs, while 5H cells were much shorter and more radial than 2H cells. The 2H and 5H cells showed sustained mRNA expression of stemness genes, such as *NANOG* and *OCT-4*, under an adipogenesis-inducing medium, although 5H cells showed lower expression than 2H cells (Fig. 1C). Additionally, the 2H and 5H cells showed up-regulation of *NANOG* and *OCT-4* only when cultured in DMEM medium (Fig. S1). 2H and 5H cells were maintained in a manner in which they retained their ability to differentiate into adipocytes in response to adipogenesis medium, despite the low potency rate compared to BMSCs (Fig. 1D). These findings suggest that 2H and 5H cells retained the characteristics of BMSCs.

### Co-overexpression of MDM2 and CDK4 synergistically drives tumorigenic phenotypes of transformed human BMSCs

To establish cells co-overexpressing MDM2 and CDK4, 2H and 5H cells were infected with either LacZ-(β-galactosidase, control) or MDM2-and/or CDK4-expresssing lentiviral particles. Expression of the transcript and protein of MDM2 and/or CDK4 was confirmed by immunoblotting (Fig. 2A) and qRT-PCR (Fig. S2). To evaluate the expression levels of MDM2 and/or CDK4 in transduced 2H and 5H cells, we compared the expression to two representative liposarcoma cell lines, LIPO-863B (WDLPS) and LP6 (DDLPS). MDM2 protein expression was increased by 1.06- and 1.37-fold and 2.63- and 4.14-fold in 2H and 5H cells, respectively, and the fold-change 2.32–3.24 in LIPO-863B cells. In addition, *MDM2* mRNA expression values were observed at biological levels, with fold-changes of 0.29 or 0.62 in 2H and 5H cells compared to in LIPO-863B cells (Fig. S2). For CDK4 expression, transduced 2H and 5H cells showed more than two-fold higher expression than in both LP6 and LIPO-863B cells (Fig. 2A). Morphologically, both 2H and 5H cells co-overexpressing MDM2 and CKD4 were much longer and thinner than those solely expressing MDM2, CDK4, or LacZ (Fig. 2B).

**Figure 2.**
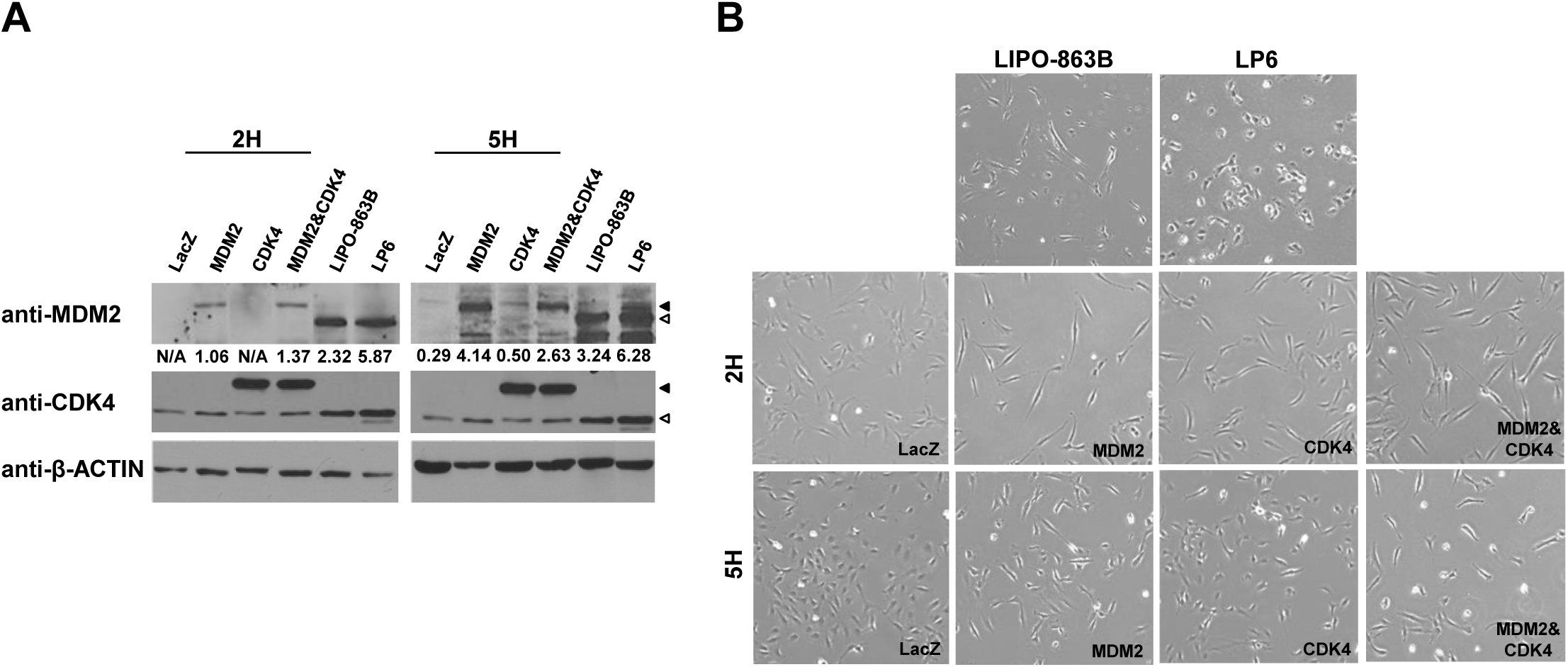
2H and 5H cells stably express MDM2 and/or CDK4 and display morphologic changes. (A) The expression of proteins of MDM2 and/or CDK4 was measured by immunoblotting. ◀, 3XFLAG-MDM2 or CDK4; ◁, MDM2 or CDK4. Fold-changes were determined by comparing the levels based on β-ACTIN using Image J. (B) Morphologic changes were assessed by comparison among different cell lines.

We next examined whether co-overexpression of MDM2 and CDK4 accelerates tumorigenic potential in transformed cells. In both 2H and 5H cells, co-overexpression of MDM2 and CDK4 significantly increased cell proliferation (Fig. 3A and 3C), anchorage-independent cell growth (Fig. 3B and 3D), and cell migration (Fig. 3E) compared to sole expression of MDM2 or CDK4. Interestingly, 5H cells only expressing MDM2 showed significantly increased anchorage-independent cell growth (Fig. 3D) and activated cell mobility (Fig. 3E), but not cell proliferation (Fig. 3C) relative to cells expressing CDK4 alone. These results indicate that co-overexpression of MDM2 and CKD4 plays a key role in tumorigenesis when transformed in BMSCs.

**Figure 3.**
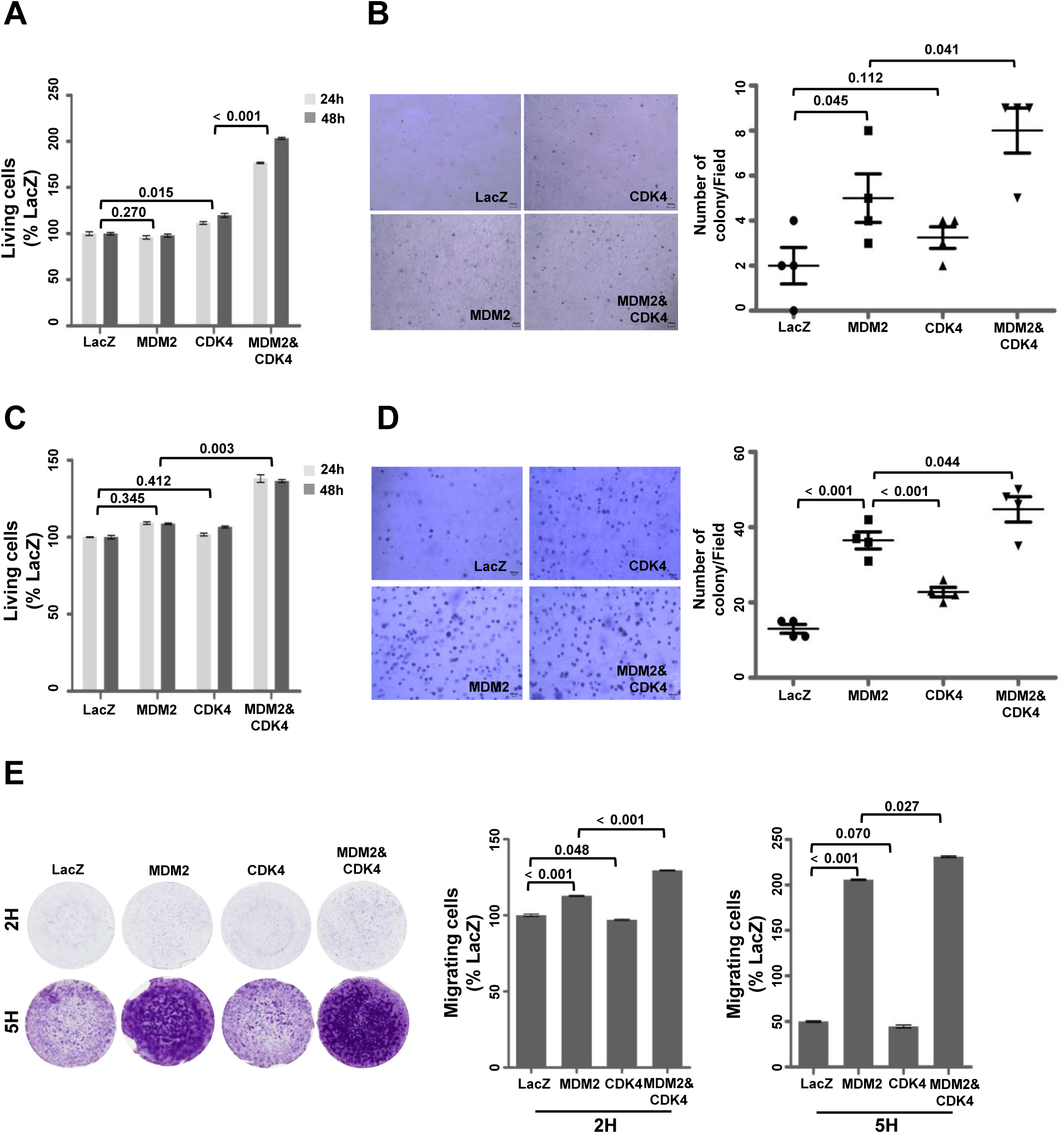
Co-overexpression of MDM2 and CDK4 synergistically promotes oncogenic potency. (A and C) Cell viability was evaluated in a WST-1 assay. (B and D) Anchorage-independent cell growth was analysed via soft agar assay. Images of cells expressing the indicated protein level in agarose are shown (left panels) and the number of colonies/fields was counted by microscopy (right panels). (E) Migration assay was performed in a Trans-well chamber. Images of crystal violet-stained cells expressing the indicated protein on membranes as shown in the left panels. Intensity was obtained by ultraviolet spectrometry and the percentage values were determined based on LacZ cell intensity (right panels). *P* values are presented for the indicated comparison.

### Co-overexpression of MDM2 and CDK4 blocks differentiation potential of adipogenesis

To examine whether overexpression of MDM2 and CDK4 alters adipogenesis potential, we first performed Oil Red O staining after culture in adipogenic induction medium. 2H-MDM2 & CDK4 cells displayed small amounts of lipid droplets compared to 2H-LacZ cells and became longer and thinner than cells expressing only MDM2 or CDK4 (Fig. 4A). 5H-MDM2 and 5H-MDM2&CDK4 cells showed reduced positivity of Oil Red O staining and contained stellar cell bodies compared to both 5H-LacZ and 5H-CDK4 cells (Fig. 4C). Next, we analysed the expression levels of genes serially induced during adipogenesis by real-time PCR: *C/EBPβ* (in the early step), *C/EBPα* and *PPARγ* (from the middle to late steps), *C/SREBP1* (full step), and *ADIPSIN* and *LPL* (late step) (Cowherd *et al*., 1999). As shown in Fig. 4B, cells expressing only MDM2 or co-overexpressing MDM2 and CDK4 showed relatively down-regulated expression levels compared to that in both 2H-LacZ and 2H-CDK4 cells. Their expression levels were similar to those of LP6 cells except for *C/EBPβ*. Compared to 5H-LacZ cells, MDM2 and/or CDK4 expression led to decreased expression of all genes except *C/EBPβ* and *PPARγ*. Notably, co-overexpression of MDM2 and CDK4 decreased the expression of all adipogenesis-related genes, showing similar levels in LP6 cells (Fig. 4D). Taken together, these data suggest that co-expression of MDM2 and CDK4 blocks the differentiation of adipogenesis from the early to late stages.

**Figure 4.**
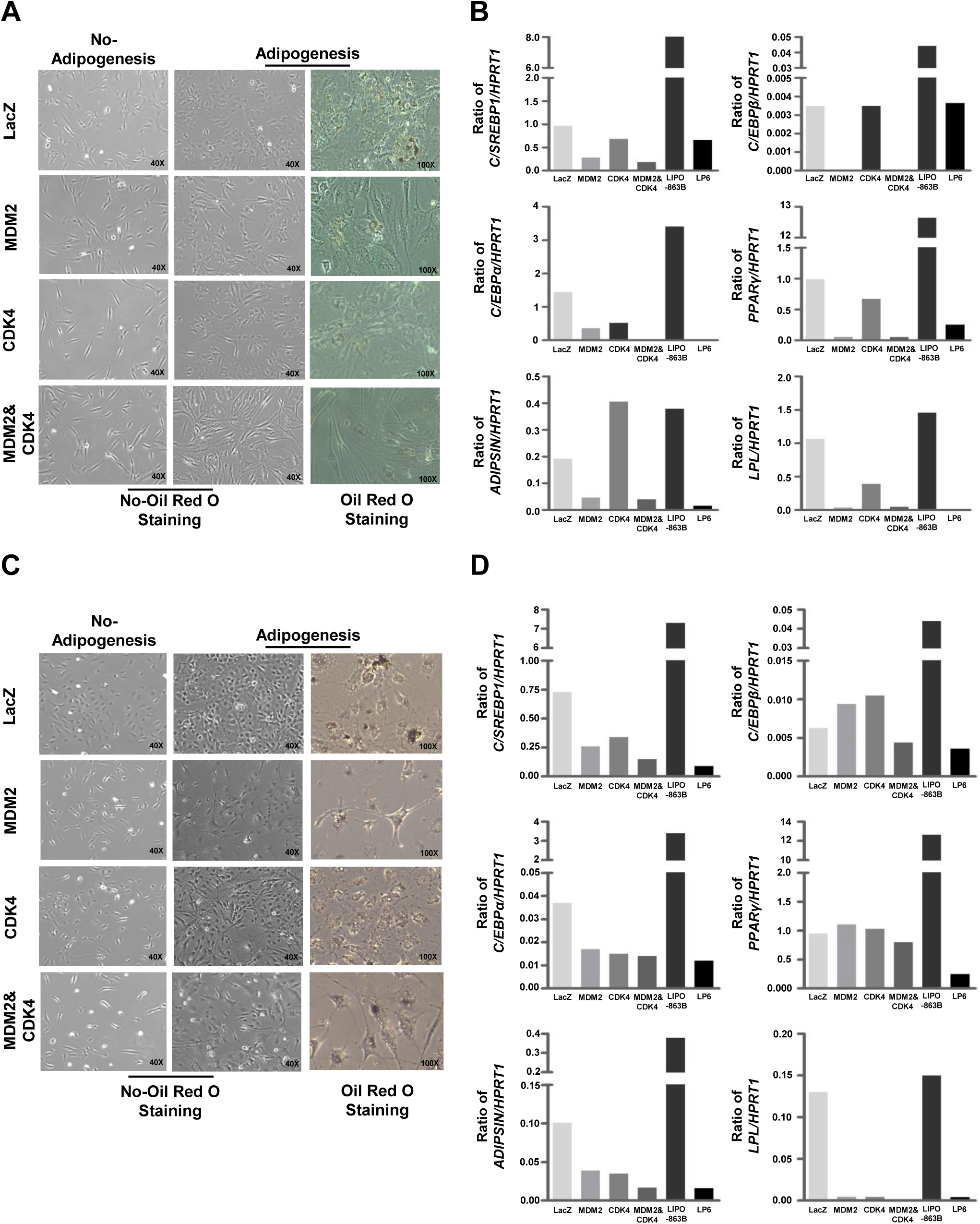
Co-overexpression of MDM2 and CDK4 blocks differentiation during full adipogenic process. (A and C) Ability to differentiate into adipocytes was evaluated by Oil Red O staining. (B and D) mRNA expression levels of *C/EBPβ*, *C/EBPα*, *PPARγ*, *C/SREBP1*, *ADIPSIN*, and *LPL* were quantified by quantitative RT-PCR. The ratio of genes to *HPRT1* was used to determine the relative levels of all genes.

### Co-overexpression of MDM2 and CDK4 in transformed human BMSCs results in the development of proliferative DDLPS tumours *in vivo*

To verify *in vivo* tumorigenic potential, nude mice were subcutaneously inoculated with 2H and 5H cells co-overexpressing MDM2 and CDK4. 2H cells did not develop into tumours, regardless of co-expression of MDM2 and CDK4 (Fig. 5A). Consistent with a study by Rodriguez *et al*., 5H-LacZ cells formed tumours with high penetrance (4/5, Fig. 5A) (Rodriguez *et al*., 2013). However, the 5H-MDM2&CDK4 cells developed larger tumours than those observed from the LacZ control (Fig. 5B and 5C). In addition, LP6 cells showed much more aggressive tumour formation compared to 5H-MDM2 and CDK4 cells *in vivo* despite their low growth potency *in vitro* (Fig. 5B and 5C, and Fig. S3).

**Figure 5.**
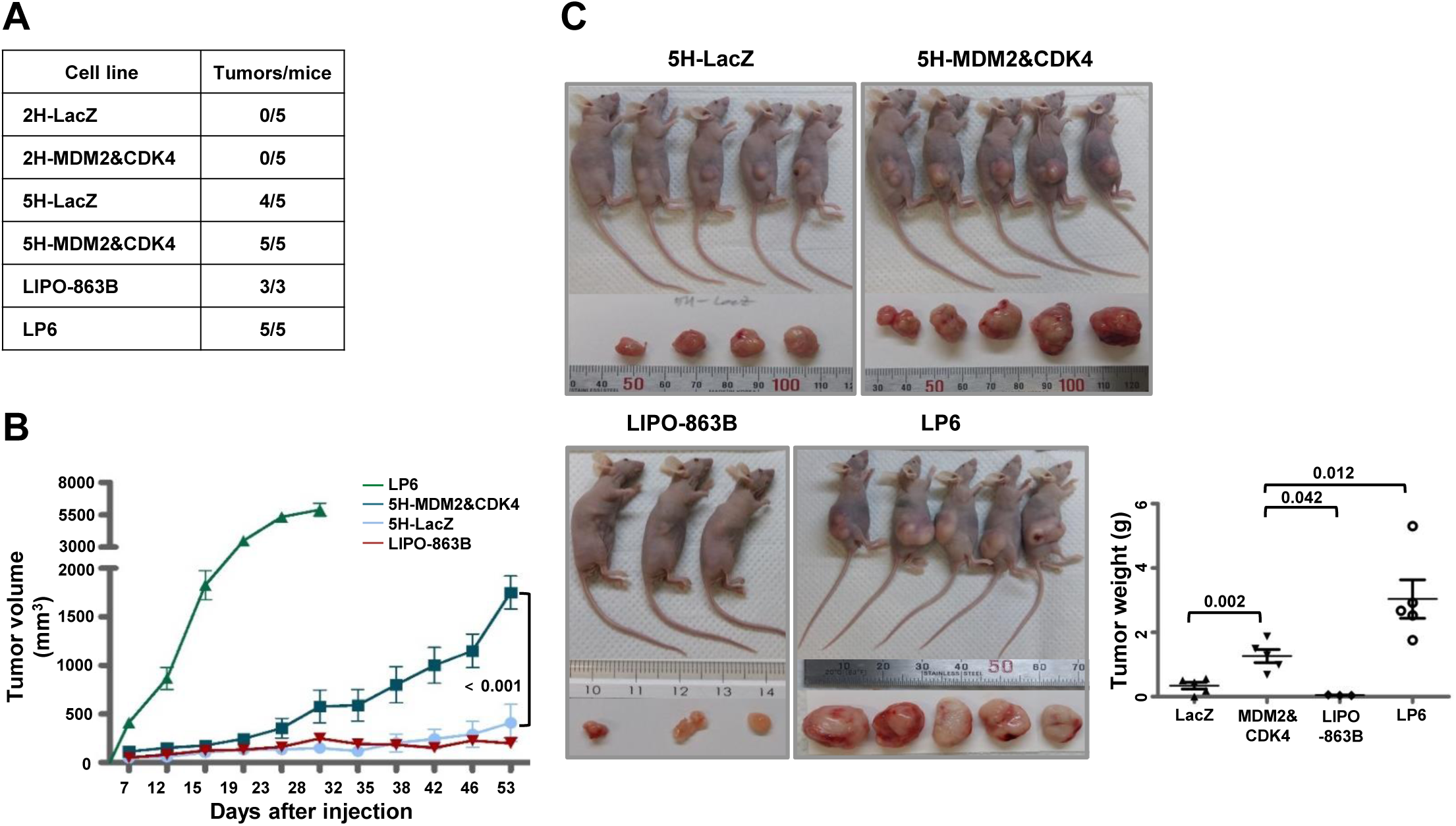
Co-expression of MDM2 and CDK4 in 5H cells aggressively causes tumour formation *in vivo*. (A) Incidence of tumour formation is illustrated. (B) Tumour volumes were measured using a digital calliper on the indicated day. (C) All xenografts were resected, and tumour weights were measured at the time of tumour harvest. *P* values are presented for the indicated comparison.

We next performed histological analysis of 5H cell-derived tumours including the LIPO-863 and LP6 xenograft models. The 5H-MDM2 and CDK4 cell-derived tumours were immunostained and found to be strongly positive for MDM2 and CDK4, while tumours derived from 5H-LacZ were not (Fig. 6A). Although relatively high background was observed, their intensity levels were similar to those of LP6 cell-derived tumours (Fig. 6A). The 5H-MDM2 and CDK4 cell-derived tumours exhibited more proliferative features with high cellularity compared to 5H-LacZ-derived tumours (Fig. 6B). Moreover, these tumours morphologically resembled LP6-derived tumours displaying large nuclear cells with variable sizes dispersed within a fibrous matrix, but not LIPO-863B-derived tumours which were composed of mature adipocytic cells with diverse size, associated with a variable number of atypical stromal cells (Fig. 6B and Fig. S4). In addition, tumours from 5H-MDM2&CDK4 cells showed a small portion of lipoblast cells but they were not immunostained with KU80 reported as a marker of human cells; therefore, these lipoblast cells may not be derived from 5H-MDM2&CDK4 cells (Fig. S5) (Allard *et al*., 2014). These findings indicate that co-overexpression of MDM2 and CDK4 in 5H cells with five additional oncogenic mutations can result in the development of proliferative DDLPS *in vivo*.

**Figure 6.**
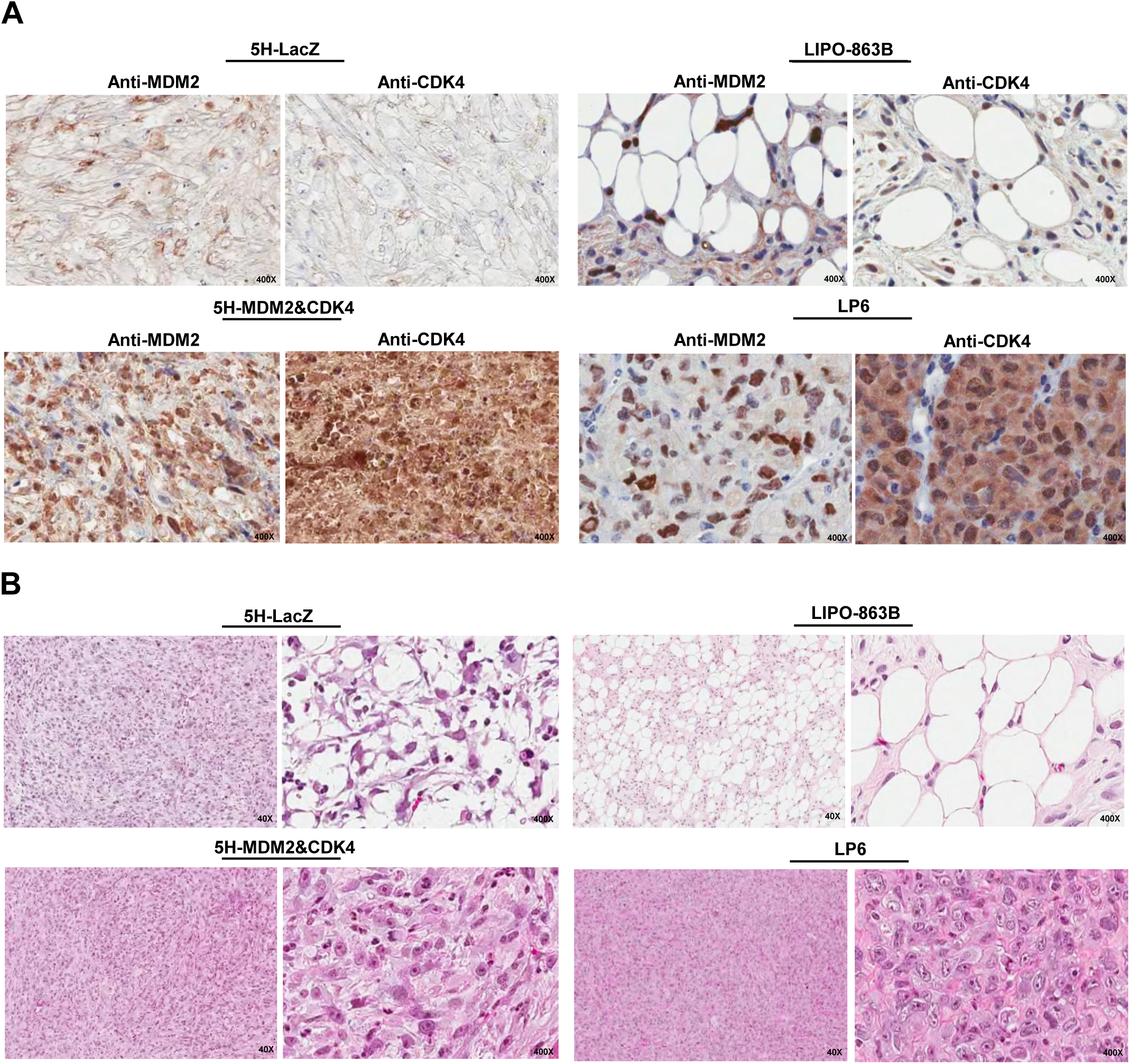
Co-expression of MDM2 and CDK4 in 5H cells induces proliferative DDLPS phenotypes *in vivo*. (A) Expression of MDM2 or CDK4 were examined by immunohistochemical staining in tumours developed from 5H-LacZ, 5H-MDM2&CDK4, LIPO-863B, and LP6 cells. Original magnification is indicated. (B) Histological characteristics of derived tumours were monitored by haematoxylin-eosin staining.

## Discussion

WDLPS/DDLPS is characterized by amplification and overexpression of MDM2 and CDK4. However, several other oncogenes, such as HMGA2, c-JUN, ZIC1, and others, have been reported to contribute to tumorigenesis and disease progression of WDLPS/DDLP (Barretina *et al*., 2010; Brill *et al*., 2010; Italiano *et al*., 2009; Italiano *et al*., 2008; Mariani *et al*., 2007; Sioletic *et al*., 2014; Snyder *et al*., 2009). Although amplification and overexpression of MDM2 and CDK4 are hallmark events of WDLPS/DDLPS, whether MDM2 and CDK4 drive WDLPS/DDLPS tumorigenesis remains unclear. Therefore, it is necessary to evaluate whether amplification and overexpression of MDM2 and CDK4 is a critical event for WDLPS/DDLPS development. We addressed this question by establishing a corresponding xenograft.

Human BMSCs have not been shown to undergo spontaneous transformation *in vitro*, other than in few cases in which BMSC-injected patients later developed osteosarcoma (Aguilar *et al*., 2007; Berger *et al*., 2008; Bernardo *et al*., 2007; Bielack *et al*., 2003). In addition, sarcomagenesis models expressing the fusion proteins EWS-FLI1 or SYT-SSX1 in human BMSCs failed to generate tumour phenotypes (Cironi *et al*., 2009; Riggi *et al*., 2008). However, genetic introduction of tumour suppressor genes, such as TP53 and RB, and other oncogenes, such as SV40 T antigen and HRAS, promoted BMSC transformation (Xiao *et al*., 2013). Rodriguez *et al*. first succeeded in inducing myxoid liposarcoma using BMSCs expressing the FUS-CHOP fusion protein and transformation with five oncogenic hits: TP53 deficiency, RB deficiency, hTERT overexpression, C-MYC stabilization, and HRAS^v12^ overexpression (Rodriguez *et al*., 2013). Thus, cooperating oncogenic hits are needed to transform BMSCs. Based on these reports, we induced WDLPS/DDLPS *in vivo* by co-overexpressing MDM2 and CDK4 in transformed BMSCs.

According to a study by Taylor *et al*., *HDAC1*, *MAPKAP1*, *PTPN9*, and *DAZAP2* were mutated in DDLPS tissue samples by next-generation sequencing analysis (Taylor *et al*., 2011). Kanojia *et al*. found that *TERT*, *MCL1*, *ROR1*, *ERBB4*, *VEGFA*, *CPM2*, *ERBB3*, *SOCS2*, *CCNE1*, and *RUNX1* were amplified, *E2F6*, *CDKN2A*, and *NF1* were deleted, and *TP53*, *PLEC*, *FAT3*, *MXRAS*, *CHEK2*, and *NF1* were significantly altered in WDLPS/DDLPS tissue samples from copy number profiling (SNP array) analysis and targeted exome sequencing (Kanojia *et al*., 2015). Notably, Ballinger *et al*. identified pathogenic germline mutations in 638 (55%) samples among 1162 sarcoma patients using a targeted exon sequencing panel comprised of 72 genes (based on associations with increased cancer risk) including *TP53* and *RB* (Ballinger *et al*., 2016). They also showed that multiple mutations are significantly correlated with poorer tumour-free survival. Here, 2H cells with two oncogenic hits, TERT and HPV-16 E6 (TP53 degradation), did not develop into tumours, while 5H cells harbouring three additional oncogenic hits, HPV-16 E7 (RB family inactivation), SV40-ST (c-MYC stabilization), and HRAS^v12^ (mitogenic signal activation), generated tumours *in vivo*. Our findings suggest that TERT and HPV-16 E6 may not cooperate with MDM2 and CDK4 for BMSCs to develop into WDLPS/DDLPS, but HPV-16 E7, SV40-ST, or HRAS^v12^ may be required to contribute to the transformation of BMSCs.

Although histological analysis of 5H-MDM2 and CDK4-derived tumours could not be clearly classified as DDLPS, co-overexpression of MDM2 and CDK4 played a key role of tumorigenesis during the transformation of BMSCs under cooperation conditions with multiple genetic alterations. DDLPS is thought to develop from WDLPS after a long duration, which was confirmed by the detection of exclusively low-grade dedifferentiated components, its specific genomic profile compared to WDLPS, and that most DDLPS occurs *de novo* (90%); therefore, DDLPS now be identified in the absence of WDLPS (Chibon *et al*., 2002; Ghadimi *et al*., 2011; Lahat *et al*., 2008). Moreover, our *in vivo* experiments showed that DDLPS tumour potency may be induced without the WDLPS component. To comprehensively understand the mechanism of WDLPS/DDLPS development, characterization of both germline and somatic genetic alterations is needed using a massive next-generation sequencing approach in different large cohorts of each WDLPS or DDLPS.

Although cell proliferation and differentiation are regarded as mutually exclusive events, cross-talk has been reported between both processes during adipogenesis (Fajas, 2003). Previous reports suggested that MDM2 promotes adipocyte differentiation through *CREB*-dependent transactivation or *CREB*-regulated transcriptional coactivator-mediated activation of STAT6 using mouse embryonic fibroblast and mouse preadipocyte cells and CDK4 participates in adipogenesis through *PPARγ* activation (Abella *et al*., 2005; Hallenborg *et al*., 2012; Hallenborg *et al*., 2016). However, Peng *et al*. showed that WDLPS/DDLPS cell lines exhibited low or negative levels of Oil Red O positivity and PPARγ compared to pre-adipocytes and adipocytes (Peng *et al*., 2011). We also found that 2H-MDM2&CDK4 and 5H-MDM2&CDK4 cells showed reduced positivity of Oil Red O staining, and co-overexpression of MDM2 and CDK4 decreased the expression of all adipogenesis-related genes. Therefore, MDM2 and/or CDK4 may function as initiating oncogenes to block adipogenic differentiation during WDLPS/DDLPS development.

In summary, co-overexpression of MDM2 and CDK4 causes DDLPS tumours in transformed human BMSCs by accelerating cell growth and migration, and blockage of adipogenetic potential under cooperation conditions with multiple genetic factors.

## Materials and Methods

### Cell lines and reagents

Transformed 2H and 5H human bone marrow stem cells and the LIPO-863B and LP6 cell lines were kindly provided by Dr. Pablo Menendez, Dina Lev, and Jonathan A Fletcher, respectively (Peng *et al*., 2011; Riggi *et al*., 2008; Sioletic *et al*., 2014). The cells were cultured in Dulbecco’s Modified Eagle Medium (DMEM) (Thermo Fisher Scientific, Waltham, MA, USA) containing 10% foetal bovine serum (FBS) (Gibco, Grand Island, NY, USA) and 1% antibiotic–antimycotic (Gibco) at 37°C and in 5% CO₂. Mycoplasma contamination was not detected in any cell.

### RT-PCR

cDNA was generated from total RNA using SuperScript III Transcriptase according to the manufacturer’s instructions (Invitrogen, Carlsbad, CA, USA). Quantitative reverse transcription (qRT)-polymerase chain reaction (PCR) amplification of stemness- or adipogenesis-related genes was conducted using probes and primers with the Universal Probe Library System (Roche, Basel, Switzerland). *MDM2* amplification was performed using the following primer pair: Forward, (5′-ACCTCACAGATTCCAGCTTCG); Reverse, (5′-TTTCATAGTATAAGT GTCTTTTT). *HPRT1* was used as a reference gene, and the ratio of each gene to *HPRT1*was calculated for relative quantification of the expression level of each gene.

### Immunoblotting, immunocytochemistry, immunohistochemistry, and fluorescence-activated cell sorting (FACS)

Equal amounts of protein were subjected to SDS-PAGE on an 8.5% gel before being transferred to a nitrocellulose membrane (Pall Corporation, Port Washington, NY, USA). The membrane was incubated with primary anti-MDM2, CDK4, and β-ACTIN (diluted 1:1,000 in 5% non-fat milk, Santa Cruz Biotechnology, Dallas, TX, USA) and FLAG (diluted 1:1,000 in 5% non-fat milk, Sigma-Aldrich, St. Louis, MO, USA) antibodies, and then washed (30 min) with T-BST. The membrane was incubated with horseradish peroxidase-conjugated secondary goat anti-rabbit or anti-mouse antibodies (diluted 1:2,000 in 5% non-fat milk, Abcam) for 1h, followed by 30 min of washing with T-BST. Signals were detected using ECL solution (Thermo Fisher Scientific). Four-micrometre-thick sections from formalin-fixed paraffin-embedded cell or tissue blocks were cut with a microtome and routinely deparaffinized. The antigen retrieval procedure was performed in 0.01 M of citrate buffer (pH 6.0) at 95°C, and counterstaining was conducted with haematoxylin. The anti-MDM2 (Invitrogen, IF2, 1:200 dilution), CDK4 (Invitrogen, DCS-31, 1:50 dilution), and KU80 (Cell Signaling, Danvers, MA, USA; C48E7, 1:200 dilution) antibodies were used for immunocytochemical or immunohistochemical staining using the automated bench-mark XT platform (Ventana Medical Systems, Tucson, AZ, USA).Cells were washed with FACS buffer (PBS, 0.5% BSA, 0.1% NaN_3_ sodium azide) and stained with anti-CD34 and CD105 (BD Biosciences, San Jose, CA, USA) antibodies. Isotype matched FITC/PE conjugated controls were also included with each set. Positive cells were analysed by BD FACSVerse flow cytometry (BD Biosciences).

### Generation of stable cell line over-expressing MDM2 and/or CDK4

The full-length cDNAs of *MDM2* or *CDK4* were generated from a cDNA library of human bone marrow stem cells. The PCR products were cloned into the N-terminal p3XFLAG-CMV-10 vector (Sigma-Aldrich). We confirmed the full sequence of wild-type *MDM2* and *CDK4* by the Sanger sequencing method. Full-length 3XFLAG-*MDM2* or 3XFLAG-*CDK4* were cloned into the gateway entry vector pCR8/GW/Topo (Invitrogen), and then subcloned into pLenti6.3/V5-DEST (Invitrogen). Full-length sequences of 3XFLAG-*MDM2* or 3XFLAG-*CDK4* were validated by Sanger sequencing. pLenti6.3/3XFLAG-*MDM2* or 3XFLAG-*CDK4* expression vectors were transfected into 293FT cells using ViraPower Packaging Mix (Invitrogen) to produce lentivirus expressing MDM2 or CDK4. After 48h, lentiviral supernatants were harvested and transduced into 2H and 5H cells in the presence of 8µg/mL of polybrene. Transduced cells were grown in DMEM complete medium for48hafterinfection, and then the medium was replaced with medium containing blasticidin (5µg/mL) after 24h. Cells were selected for 2 weeks using selective medium. Stable expression of MDM2 or CDK4 was confirmed by qRT-PCR and immunoblotting.

### Cell proliferation and migration assays and soft agar assay

The cell proliferation assay was performed using an EZ-CYTOX kit (Daeil Lab Service, Seoul, Korea) according to the manufacturer’s instructions. Cells were plated in 96-well plates (3 ×10^2^cells/well). The 96-well plates were incubated with EZ-CYTOX reagent for 3 hours at 37°C after 1 and 2 days. Twenty-four-well trans-well chambers (Corning Costar, Corning, NY, USA) with 8-μm polycarbonate membrane filters were used to determine migration ability. For this assay, 5 × 10^4^ cells were seeded into the upper chamber in DMEM without FBS. The lower chamber contained 700 μL of DMEM containing 10% FBS. The trans-well chamber was incubated at 37°C in 5% CO_2_. After 24 h of incubation, non-migrating cells on the upper filter surface were removed with a cotton swab and the migrated cells were stained with 0.5% crystal violet. Cells were seeded into 24-well plates with the appropriate concentrations of agarose (0.5% for base and 0.3% for top) to form colonies in 3 weeks. Colonies were stained with crystal violet (0.5% w/v) and counted using a microscope.

## Adipogenic differentiation

BMSCs and 2H and 5H cells were seeded onto a 6-well plate in DMEM medium, and then replaced with adipogenic differentiation medium (StemPro Adipogenic Differentiation Kit, Invitrogen) every 3–4 days. After 10 days, the cells were stained with an Oil Red O staining kit (Lifeline, Frederick, MD, USA) according to the manufacturer’s instructions.

## Mouse xenograft modelling

This study was reviewed and approved by the Institutional Animal Care and Use Committee of Samsung Biomedical Research Institute (SBRI, Seoul, Korea). SBRI is an Association for Assessment and Accreditation of Laboratory Animal Care International accredited facility and abides by the Institute of Laboratory Animal Resources guide (No. 20160108001). Female nude mice were injected subcutaneously with 2H, 5H, LIPO-863B, or LP6 (5×10^6^) cells. After the indicated number of days, tumour diameter was measured using a digital calliper two or three times per week, and tumour sizes were estimated using the following formula: (3.14/6) (length×width^2^).

## Acknowledgments

We would like to thank Dr. Pablo Menendez, Dr. Dina Lev, and Dr. Jonathan A Fletcher for their generous gifts and Dr. Je-Gun Joung (Samsung Genome Institute, Samsung Medical Center) for valuable advice, Dr. Sun Myeong Lee (Research Institute of Pharmaceutical Science, College of Pharmacy, Seoul National University) for technical supports. We thank Ka-Won Noh for English proofreading. This work was supported by a National Research Foundation of Korea (NRF) grant funded by the Korean Government (grant number 2013R1A1A2011536 and 2016R1A5A2945889).

## Author contributions

Yu Jin Kim and Yoon-La Choi designed the study. Yu Jin Kim and Yoon-La Choi wrote the manuscript. Yoon-La Choi reviewed the histopathology and immunohistochemistry results. Yu Jin Kim, Minjung Sung, Dan Bi Yu, Mingi Kim, Ji-Young Song, Kyoung Song, and Kyungsoo Jung performed the experiments. Yu Jin Kim and Yoon-La Choi provided critical comments on the manuscript.

---

**Supplementary Figure 1.** mRNA expression levels of *NANOG* and *OCT-4* were quantified by quantitative RT-PCR in DMEM medium. The ratio of genes to *HPRT1* was used to determine the relative levels of all genes. Percentage values were calculated based on their levels in BMSCs.

**Supplementary Figure 2.** mRNA expression of *MDM2* was measured by quantitative RT-PCR. Fold-changes were determined by comparing the levels expressed in LacZ and LIPO-863B cells.

**Supplementary Figure 3.** Cell viability was evaluated in a WST-1 assay. *P* values are presented for the indicated comparison.

**Supplementary Figure 4.** STR profile was confirmed through dissection in lipoblasts from LIPO-863B cell-derived tumour.

**Supplementary Figure 5.** Expression of KU80 was examined by immunohistochemical staining in lipoblasts from 5H-MDM2&CDK4 cell-derived tumours.

